# The temporal transcription factor E93 controls dynamic enhancer activity and chromatin accessibility during development

**DOI:** 10.1101/679753

**Authors:** Spencer L. Nystrom, Matthew J. Niederhuber, Daniel J. McKay

**Affiliations:** Curriculum in Genetics and Molecular Biology, The University of North Carolina at Chapel Hill; Department of Biology, The University of North Carolina at Chapel Hill; Department of Genetics, The University of North Carolina at Chapel Hill; Integrative Program for Biological and Genome Sciences, The University of North Carolina at Chapel Hill

**Author notes:** equal contributors (Spencer L. Nystrom, Matthew J. Niederhuber).

## Abstract

How temporal cues combine with spatial inputs to control gene expression during development is poorly understood. Here, we test the hypothesis that the *Drosophila* transcription factor E93 controls temporal gene expression by regulating chromatin accessibility. Precocious expression of E93 early in wing development reveals that it can simultaneously activate and deactivate different target enhancers. Notably, the precocious patterns of enhancer activity resemble the wild-type patterns that occur later in development, suggesting that provision of E93 alters the competence of enhancers to respond to spatial cues. Genomic profiling reveals that precocious E93 expression is sufficient to regulate chromatin accessibility at a subset of its targets. These accessibility changes mimic those that normally occur later in development, indicating that precocious E93 accelerates the wild-type developmental program. Further, we find that target enhancers that do not respond to precocious E93 in early wings become responsive after a developmental transition, suggesting that parallel temporal pathways work alongside E93. These findings support a model wherein E93 expression functions as an instructive cue that defines a broad window of developmental time through control of chromatin accessibility.

## INTRODUCTION

*Cis*-regulatory regions such as enhancers and promoters interpret multiple types of inputs to control gene expression during development. These inputs come in the form of both spatial and temporal cues. Spatial cues are often provided by transcription factors, which can contribute information on cell identity (e.g. MyoD), organ identity, (e.g. Pha-4), and regional identity (e.g. Hox factors). Additional spatial cues are provided by the activity of signaling pathways such as the Wnt, BMP, and EGFR families, which contribute information on distance relative to the source of the signal through their downstream transcriptional effectors. Remarkably, many of these spatial cues are used reiteratively over the course of development, often with different effects on target gene expression. For example, the Hox factor Ubx controls different sets of target genes at different times in *Drosophila* appendage development, as does the intestine-specifying factor CDX2 during gut development in mouse and humans (Kumar et al., 2019; Pavlopoulos and Akam, 2011). Similarly, EGFR signaling promotes wing vein formation early in *Drosophila* larval development, whereas later in pupal stages, EGFR represses vein formation and instead promotes differentiation of the complementary intervein cells (Martín-Blanco et al., 1999). Thus, spatial inputs alone are insufficient to account for the sequence of gene expression and cell state changes that occur during development.

Temporal cues provide an additional axis of information that can increase the range of gene expression responses to spatial inputs. Some temporal cues come in the form of post-transcriptional regulators, such as lin-4, lin-28, and let-7 in *C. elegans*, which control transitions between developmental stages through regulation of RNA stability and translation efficiency (Pasquinelli and Ruvkun, 2002). Other temporal cues come in the form of developmentally-restricted expression of transcription factors. For example, in mammals and in *Drosophila*, the diversity of cell types found in the adult nervous system depends on a temporal cascade of transcription factor expression in neural progenitor cells (Doe, 2017; Holguera and Desplan, 2018). Yet another means of temporal gene regulation involves systemically-secreted signals that coordinate the timing of gene expression programs between distant tissues. Ecdysone signaling in insects and thyroid hormone-dependent metamorphosis in amphibians are classic examples of systemic signals that trigger temporal-specific gene expression changes during development.

Although it is clear that both spatial and temporal inputs are necessary for proper gene regulation during development, the mechanisms by which these inputs combine to control target enhancer activity are poorly understood. One potential mechanism for control of the responsiveness of enhancers to transcriptional inputs is regulation of chromatin accessibility. DNA that is wrapped around histone proteins to form nucleosomes, also referred to as “closed” chromatin, is typically refractory to transcription factor binding. By contrast, depletion or remodeling of nucleosomes, such that an enhancer is made “open” or “accessible,” creates a permissive site for transcription factors to bind DNA. In principle, the accessibility of an enhancer could determine whether it is competent to respond to transcription factor input and thereby help to explain how they can be reutilized during development with different transcriptional outcomes.

Recently, support for the role of chromatin accessibility in the integration of spatial and temporal factor inputs has emerged from examination of tissues at two different stages of development in *Drosophila*: neural diversification in the embryo and specification of appendage cell fates in the pupa (McKay and Lieb, 2013; Sen et al., 2019; Uyehara et al., 2017). In the embryo, distinct neural stem cell lineages are determined by differential expression of spatial transcription factors. Within a given lineage, neural stem cells utilize sequential expression of temporal transcription factors to produce progeny with distinct identities over time. Importantly, different neural lineages use the same set of temporal transcription factors to specify progeny identities. Using a lineage-specific method of generating genome-wide DNA binding profiles, the temporal transcription factor Hunchback was found to bind different target sites in different neural lineages. Moreover, these target sites correspond to lineage-specific patterns of open chromatin (Sen et al., 2019). These findings indicate that the temporal factor Hunchback does not control open chromatin profiles and does not determine where it binds in the genome. Instead, they suggest that the spatial transcription factors expressed in neuroblasts control open chromatin profiles to drive lineage-specific binding of temporal transcription factors.

The ecdysone-induced transcription factor E93 provides a contrasting example of temporal transcription factor function. Similar to Hunchback’s role in the embryonic nervous system, E93 functions as a temporal identity factor. E93 is activated during the transition from prepupal to pupal stages of metamorphosis, and E93 loss-of-function mutations exhibit defects in cell fates that are specified during this time (Baehrecke and Thummel, 1995; Mou et al., 2012). Also similar to Hunchback, E93 combines with spatial cues to pattern cell fates. During specification of the pigmented bract cells, which are induced adjacent to bristles during pupal leg development, E93 expression makes the *Distal-less* gene competent to respond to EGFR signaling (Mou et al., 2012). However, in contrast to Hunchback, recent work from pupal wings suggest a direct role for E93 in regulating chromatin accessibility (Uyehara et al., 2017). During metamorphosis, the wing imaginal disc undergoes dramatic morphological, cell fate, and gene expression changes to form the notum (back), hinge, and wing blade of the adult. Gene expression and chromatin accessibility profiling of larval wing imaginal discs and pupal wings 24-hours and 44-hours after puparium formation revealed that these changes coincide with sequential changes in open chromatin genome-wide (Guo et al., 2016; Uyehara et al., 2017). These changes in chromatin accessibility are strongly correlated with enhancer activity. Sites that open with time correspond to late-acting enhancers switching on, and sites that close with time correspond to early-acting enhancers switching off. Many of these temporally-dynamic open chromatin sites are bound by E93 in pupal wings. Moreover, chromatin accessibility profiling of E93 mutants determined that E93 is required for temporal changes in chromatin accessibility and enhancer activity. In the absence of E93, early-acting target enhancers fail to close and fail to turn off. Conversely, late-acting target enhancers fail to open and fail to turn on (**Figure 1A**). These findings suggest that E93 is directly required for many of the sequential changes in chromatin accessibility that occur in pupal wing development. More generally, they support a model in which E93 functions as a temporal identity factor by acting as a gatekeeper that controls access to *cis*-regulatory elements in the genome. In this model, E93 acts as a temporal identity factor by making late-acting enhancers competent to respond to spatial inputs by increasing their accessibility, whereas it deactivates early-acting enhancers by decreasing their accessibility, thus allowing for reutilization of spatial inputs and driving development forward in time.

**Fig. 1.**
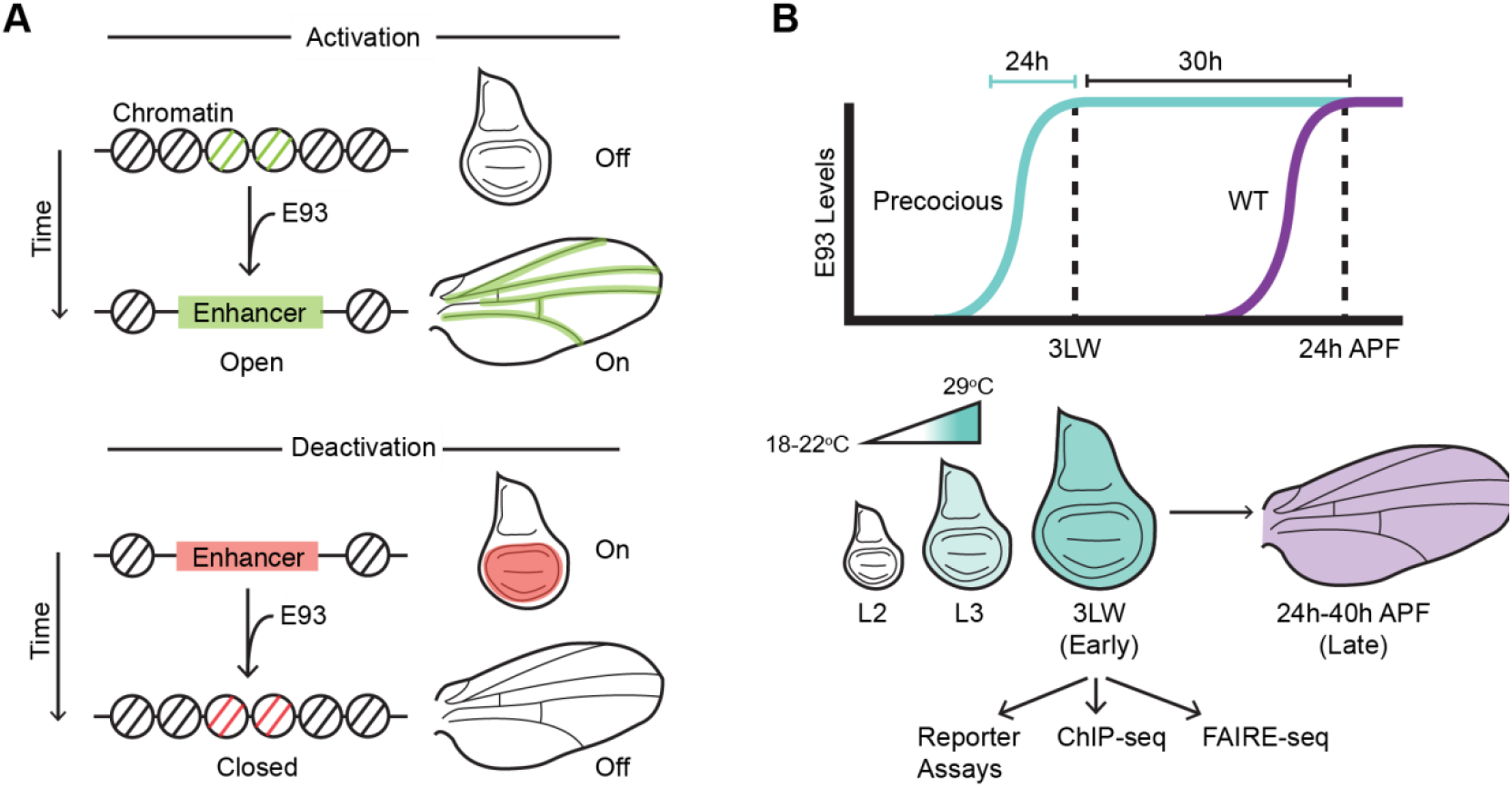
Illustrations of E93-mediated enhancer regulation and experimental design. (A) Schematic depicting proposed role of E93 in opening and activating late-acting enhancers (top) and closing and deactivating early-acting enhancers (bottom). Patterns of enhancer activity are indicated in green and red. (B) Schematics depicting the timing of wild-type E93 expression (purple) relative to the timing of precocious expression experiments (teal). GAL4 drivers in combination with GAL80^ts^ were used to precociously express E93 in third instar wings (L3) for subsequent dissection at third larval wandering (3LW).

In this study, we sought to address two major unanswered questions regarding E93-dependent control of enhancer activity. First, E93 appears to simultaneously coordinate the opening and activation of certain target enhancers while closing and deactivating others, however, the determinants of this context-specific activity are unknown. Second, we sought to determine the sufficiency of E93 expression to regulate target enhancers. Although E93 is required for sequential changes in chromatin accessibility, it is not known if E93 simply maintains accessibility changes initiated by other factors, or if it initiates these changes itself.

We took advantage of the temporal sequence of *Drosophila* wing development to investigate the limits of E93 function by expressing it at an early stage of wing development, prior to the endogenous E93 expression window. We find that precocious E93 expression is sufficient to regulate target enhancer activity, and that it can simultaneously activate and deactivate different target enhancers in the same cells. Genome-wide profiling demonstrates these findings are generalizable, and that precocious E93 expression accelerates the wild-type chromatin accessibility program. Finally, we find that not all E93 target enhancers respond to precocious E93 expression in wing imaginal discs, even after prolonged exposure to E93. However, these target enhancers become responsive to precocious E93 later in prepupal wings, suggesting the requirement of an additional temporal input that is independent of E93. Together, this work supports a model in which E93 expression defines a broad temporal window, providing competence of genes to respond to inductive signals by regulating chromatin accessibility at target enhancers.

## RESULTS

Prior work indicates that E93 controls the competence of genes to respond to developmental cues. We hypothesize a mechanism by which E93 functions as a competence factor is by controlling chromatin accessibility at target enhancers. To help define the limits of E93’s abilities to regulate enhancer activity and chromatin accessibility we designed an experimental system to express E93 outside of its normal developmental context. In wild-type animals E93 expression is temporally-regulated. During early stages of wing development, including third instar larvae, E93 is absent. It is not until later during pupal stages that ecdysone signaling directly induces E93, with transcript levels peaking by 24-hours after the larval-to-pupal transition (24hr after puparium formation, 24hAPF) (**Figure 1B,** Uyehara et al., 2017). Using *GAL4* drivers in combination with a *UAS-E93* transgene and a ubiquitously-expressed temperature-sensitive GAL80 repressor (GAL80^ts^), we induced E93 in the wing imaginal disc of third instar wandering larvae (3^rd^ Larval Wandering, 3LW), prior to when E93 is normally expressed (**Figure 1B, S1,** Bischof et al., 2013). By switching between the permissive and restrictive temperatures for GAL80^ts^, we limited the duration of exogenous E93 expression to 15–24-hours at the end of larval development. We refer to this as “precocious” E93 expression. Immunofluorescence experiments with E93 antibodies indicated that precocious E93 levels in 3LW wing imaginal discs are approximately two-fold greater than endogenous E93 levels in pupal wings (**Figure S1**). Thus, this experimental design allows us to determine the impact of near-physiological levels of E93 on enhancer activity and chromatin accessibility.

### Precocious E93 expression is sufficient to deactivate a target enhancer

We first examined whether precocious expression of E93 is sufficient to regulate target enhancer activity. Previously, we identified an E93-bound enhancer termed *br*^*disc*^ that depends on E93 for proper repression in pupal wings. In wild-type larvae, *br*^*disc*^ is active throughout the wing imaginal disc epithelium (**Figure 2A**). Later, in 30-hour pupae, *br*^*disc*^ has turned off (**Figure 2A**). In E93 mutant wings, the *br*^*disc*^ enhancer fails to turn off (Uyehara et al., 2017). To test the impact of precocious E93 expression on *br*^*disc*^ activity, we expressed E93 in the anterior-half of the wing imaginal disc with *Ci-GAL4*. We found that *br*^*disc*^ activity is strongly reduced in E93-expressing cells relative to the wild-type posterior-half of 3LW discs (**Figure 2B**). We reasoned that E93-dependent repression of *br*^*disc*^ could result either from E93 blocking the initial activation of *br*^*disc*^, or from E93 turning off *br*^*disc*^ after its initial activation. To discriminate between these possibilities, we assessed enhancer activity after only 5-hours of E93 induction at 29°C, prior to 3LW. We observed *br*^*disc*^ activity across the wing disc with no appreciable change in reporter activity in E93-expressing cells relative to E93-nonexpressing cells (**Figure 2B**). Thus, precocious E93 deactivates *br*^*disc*^ instead of simply blocking its activation. We observed similar deactivation of *br*^*disc*^ using *En-GAL4* to precociously express E93 in the posterior wing compartment (**Figure S2**). Together, these findings demonstrate that precocious E93 expression is sufficient to deactivate the *br*^*disc*^ enhancer, consistent with the requirement for E93 in turning off this enhancer in pupal wings.

**Fig. 2.**
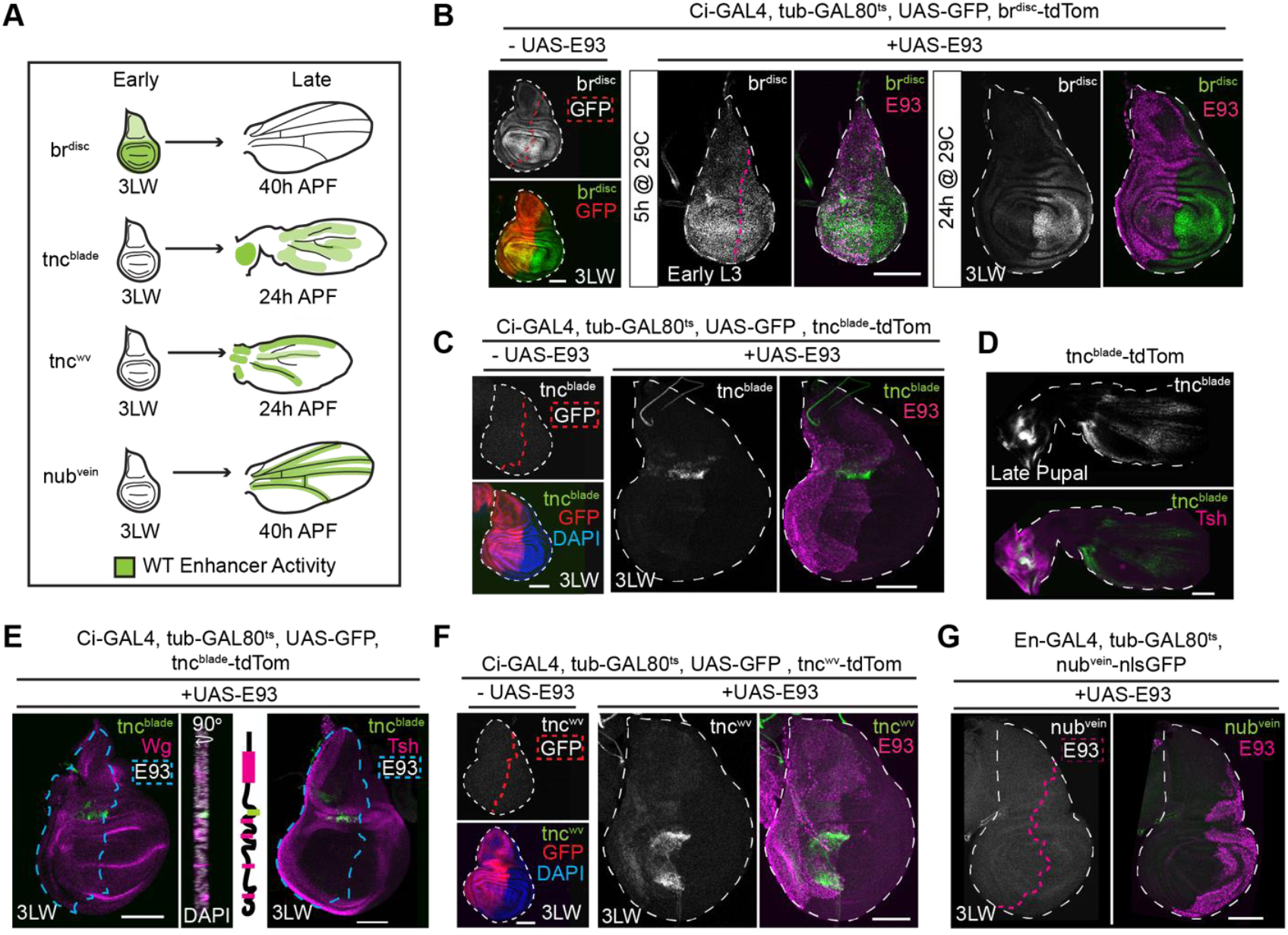
Precocious E93 expression is sufficient to deactivate and activate target enhancers. (A) Illustration depicting the WT patterns and timing of four E93-dependent enhancers (Uyehara et al, 2017). (B) Confocal images of *br*^*disc*^ activity (green) in wings. Control wings do not contain the *UAS-E93* transgene (GFP in red, left panels). Wings expressing E93 (magenta) 5-hours after induction (early L3, center panels) and 24-hours after induction (3LW, right panels) are shown. Confocal images of *tnc*^*blade*^ activity in wings (green). Control wings do not contain the *UAS-E93* transgene (GFP in red, left panels). Wings expressing E93 (magenta) 24-hours after induction are shown (right panels). (D) Confocal image of wild-type pupal wing stained for Tsh (magenta) and *tnc*^*blade*^ (green). (E) Confocal images of *tnc*^*blade*^ activity (green) in wings precociously-expressing E93 (not shown) co-stained for Wg (magenta, left panel) or Tsh (magenta, right panel). A sagittal view is also shown for the Wg-stained disc on the left. (F) Confocal images of *tnc*^*blade*^ (green) in control wings (left panels) and wings precociously-expressing E93 (magenta, right panels). (G) Confocal images of *nub*^*vein*^ (green) in wings precociously-expressing E93 (magenta). Scale bars = 100µm

### Precocious E93 expression is sufficient to activate target enhancers

In addition to *br*^*disc*^, we examined two E93-bound enhancers from the *tenectin* (*tnc*) locus that depend on E93 for proper activation in pupal wings (Uyehara et al., 2017). *Tnc* encodes a constituent of the extracellular matrix that binds integrins (Fraichard et al., 2010). In wild-type flies, the *tnc*^*blade*^ enhancer is inactive in larval wing imaginal discs. Later in 24-hour pupae, *tnc*^*blade*^ is active in the tissue between the longitudinal veins of the wing blade and at high levels within the proximal wing hinge as determined by staining late pupal wings with the proximal patterning factor Teashirt (Tsh) (**Figure 2A,D**, Zirin and Mann, 2007). Precocious expression of E93 with *Ci-GAL4* resulted in precocious activation of *tnc*^*blade*^ in 3LW wing discs in a small stripe of cells proximal to the pouch, running along the anterior-posterior (AP) axis (**Figure 2C**). Cells in which E93 precociously activates *tnc*^*blade*^ are proximal to the outer ring of Wingless (Wg), which marks the distal edge of the hinge in larval wing discs, and they are within the Tsh-expressing domain, which marks the proximal hinge (**Figure 2E**, Zirin and Mann, 2007). *Tnc*^*blade*^ was also activated in the posterior wing compartment upon precocious E93 expression using *En-GAL4* (**Figure S2**). Thus, precocious E93 expression activates the *tnc*^*blade*^ enhancer in the presumptive proximal hinge of wing imaginal discs. Notably, this pattern of precocious *tnc*^*blade*^ activation is similar to its wild-type pattern in the hinge of 30-hour pupal wings.

We observed similar outcomes with a second enhancer from the *tnc* locus, the *tnc*^*wv*^ enhancer. In wild-type wings, *tnc*^*wv*^ is active in 10-20 cells surrounding veins in 24-hour pupal wings, and like *tnc*^*blade*^, it is also dependent on E93 for activation (**Figure 2A**) (Uyehara et al., 2017). Precocious expression of E93 with *Ci-GAL4* resulted in precocious activation of *tnc*^*wv*^ in E93-expressing cells flanking the dorsal-ventral (DV) boundary, adjacent to the AP boundary in the pouch of 3LW wing imaginal discs (**Figure 2F**). Using *En-GAL4* to drive precocious E93 expression in the posterior compartment resulted in a similar pattern of precocious *tnc*^*wv*^ activity (**Figure S2**). With both *GAL4* drivers, *tnc*^*wv*^ was activated most strongly in cells within the pouch adjacent to the AP boundary, with modestly higher levels of activation further from the DV boundary (**Figure S2**). Precocious activation of *tnc*^*wv*^ near to the AP boundary in both the anterior and posterior compartment indicates the enhancer may respond to a spatial signal such as Decapentaplegic (Dpp), which is expressed in a stripe of anterior cells along the posterior compartment in wing imaginal discs. Later in pupal wings, Dpp expression decays along the AP boundary and instead becomes expressed in the presumptive veins, where it is required for vein cell fates (de Celis, 1997). Extracellular matrix constituents are among the set of Dpp targets in pupal wing veins (O’Keefe et al., 2014). Thus, similar to our observations with the *br*^*disc*^ enhancer, these findings demonstrate that precocious E93 expression is sufficient to precociously switch on late-acting pupal wing enhancers. Moreover, the pattern of precocious enhancer activity appears to be guided by similar spatial signals as the wild-type pattern of activity observed later in development, consistent with the proposed role of E93 as a competence factor.

### Precocious E93 expression is not sufficient for activation of the *nub^vein^*enhancer

Our observations with the *br* and *tnc* enhancers suggest that precocious expression of E93 accelerates the developmental program active in pupal wings. However, not all E93 target enhancers are sensitive to precocious E93 expression. We previously identified the E93-bound *nub*^*vein*^ enhancer, which is normally inactive in early wings and becomes active in 24-hour pupal wings in an E93-dependent manner (**Figure 2A**, Uyehara et al., 2017). However, in contrast to the *tnc* enhancers, precocious expression of E93 with *En-GAL4* does not activate *nub*^*vein*^ in wing imaginal discs (**Figure 2G**). We observed no difference in reporter activity in E93-expressing cells relative to their wild-type counterparts in the anterior compartment. Thus, for a subset of target enhancers, E93 expression can support activation outside of their normal developmental context. However, other target enhancers require regulatory inputs in addition to E93 for precocious activation.

### Precocious E93 binds late targets

To expand our understanding of E93’s ability to regulate target enhancers outside of its normal developmental context, we next performed a series of genome-wide profiling experiments in which E93 was precociously expressed throughout the 3LW wing imaginal disc (**Figure S1**). We first sought to define the DNA binding profiles of precocious E93 by performing ChIP-seq against HA-tagged precocious E93. Comparison of ChIP-seq profiles for precocious E93 in early wings and endogenous E93 in 24-hour pupal wings (late E93 targets) revealed three distinct binding site categories: precocious E93 binding sites that overlap late E93 targets (entopic sites), precocious E93 binding sites that do not overlap late E93 targets (ectopic sites), and late E93 binding sites that do not overlap precocious E93 targets (orphan sites) (**Figure 3A**). 82% of endogenous late targets are bound by precocious E93, suggesting that the ability of E93 to recognize and bind most of its target sites is not dependent on a late-wing developmental context (**Figure 3B**). Notably, the *br*^*disc*^, *tnc*^*blade*^, and *tnc*^*wv*^ enhancers are all bound by precocious E93, consistent with their responsiveness in reporter assays (**Figure 4D-E**). By contrast, the *nub*^*vein*^ enhancer exhibits low-level binding of precocious E93, indicating that its failure to activate in response to precocious E93 may be due to the inability of E93 to bind *nub*^*vein*^ in early wings (**Figure 5E**).

**Figure 3.**
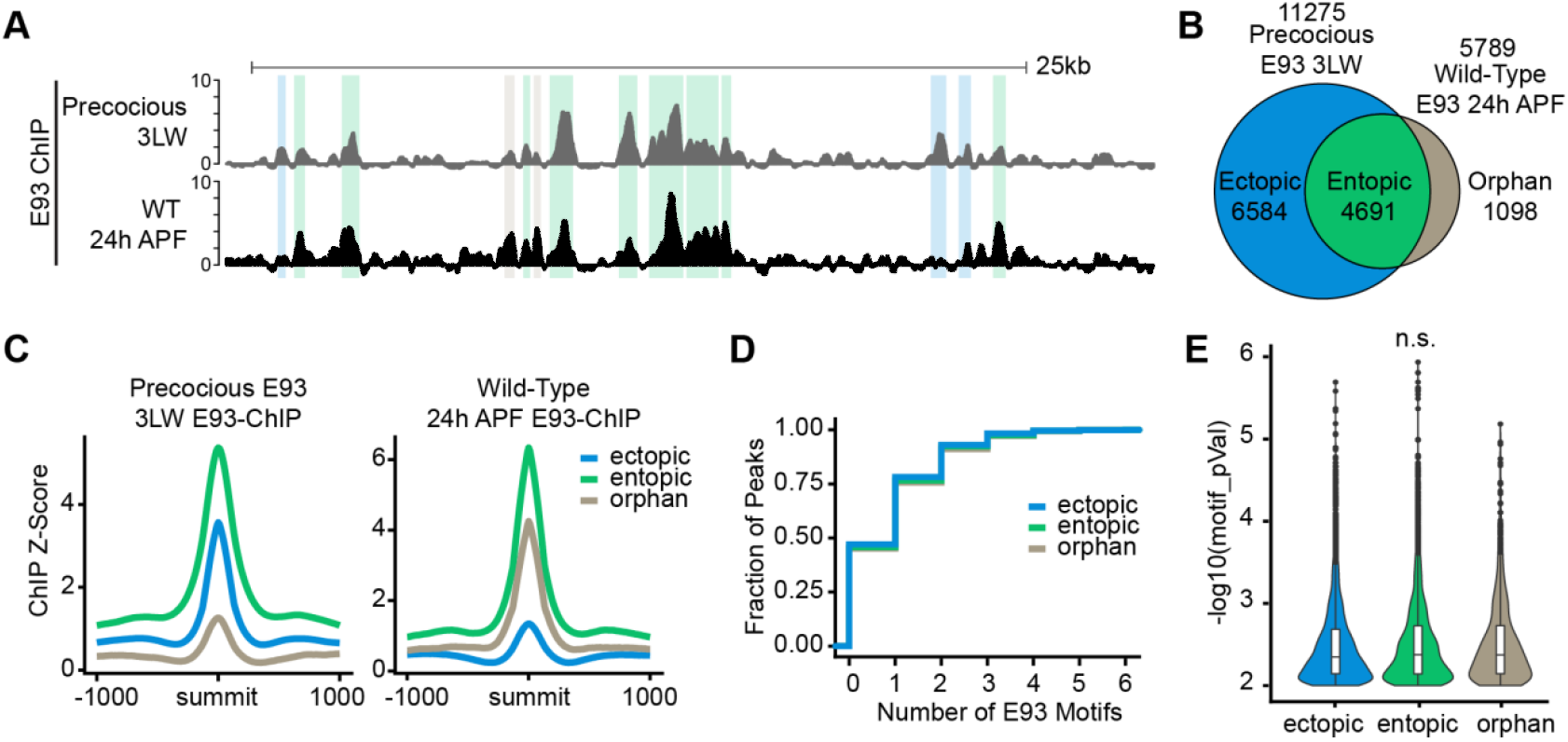
Precocious E93 recapitulates wild-type late profile in addition to binding ectopic sites. (A) Browser shot of ChIP-seq data for WT and precocious E93 wings. Colored highlights correspond to ectopic (blue), entopic (green), and orphan (brown) sites. (B) Venn diagram of peak overlaps between WT ChIP and precocious E93 ChIP-seq datasets. (C) Average signal plots of ChIP-seq Z-score within each binding category for precocious E93 and WT E93 ChIP. Cumulative distribution of the number of E93 motifs within 20bp of the summit for each binding category. (E) Violin plots depicting motif quality (–log_10_ p-value) for all E93 motifs within 20 base-pairs of E93 ChIP peak summits for each binding category (p > 0.05, one-way analysis of means, not assuming equal variance).

**Fig. 4.**
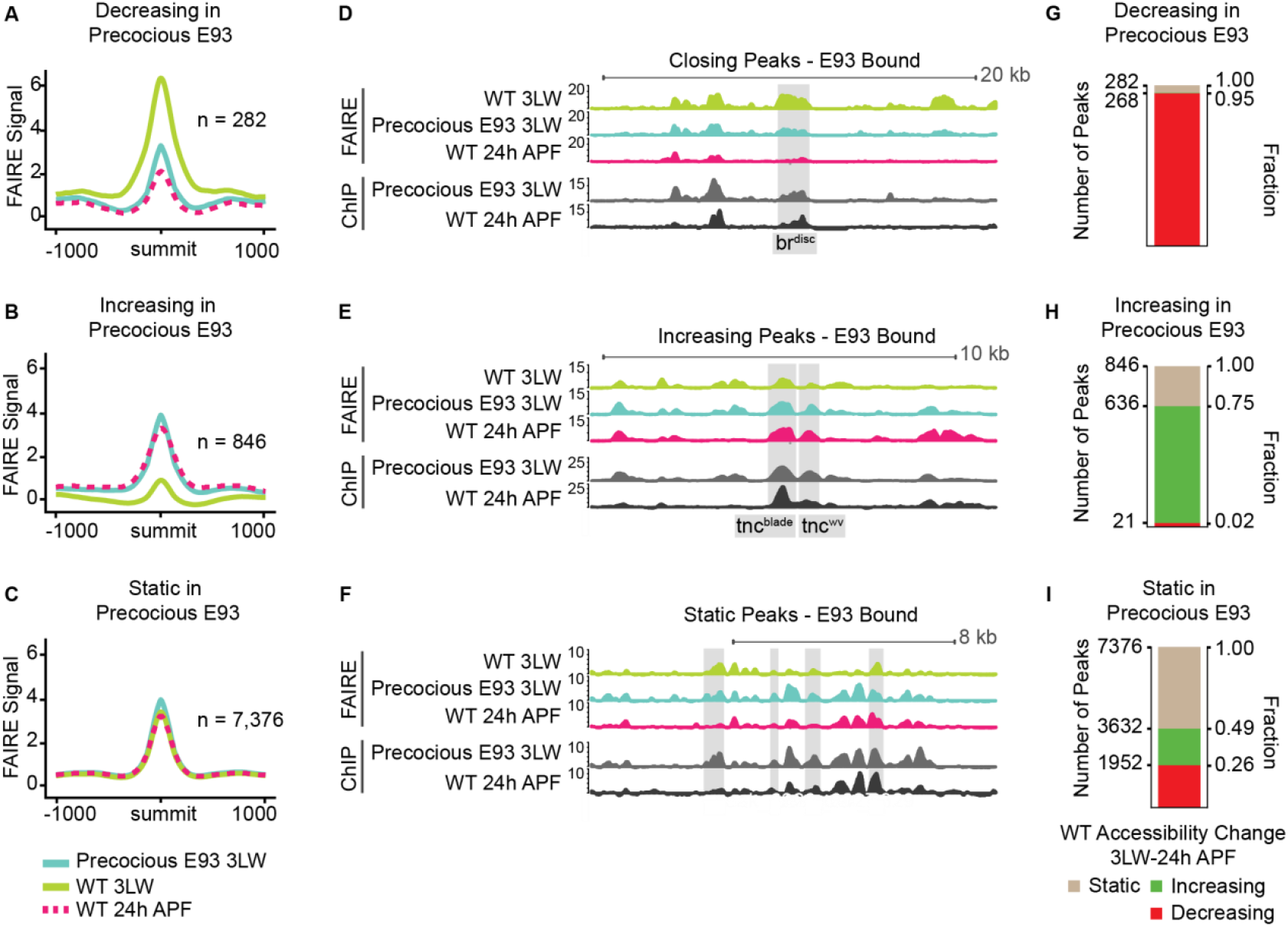
Precocious E93 accelerates the wild-type chromatin accessibility program. (A–C) Average FAIRE-seq z-scores from WT 3LW wings (green), WT 24hAPF wings (dashed red), and precocious E93 3LW wings (teal) at precocious E93 binding sites that decrease accessibility (A), increase in accessibility (B), or remain static (C) in response to precocious E93 expression. (D) Browser shot of FAIRE-seq and ChIP-seq signal (z-scores) at the *br*^*disc*^ enhancer (shaded region). Browser shot at the *tnc*^*wv*^ and *tnc*^*blade*^ enhancers (shaded regions). (F) Browser shot of static sites bound by precocious E93. (G-I) Stacked bar charts indicating the changes in chromatin accessibility that occur in wild-type development for each of the three E93 binding site categories.

**Fig. 5.**
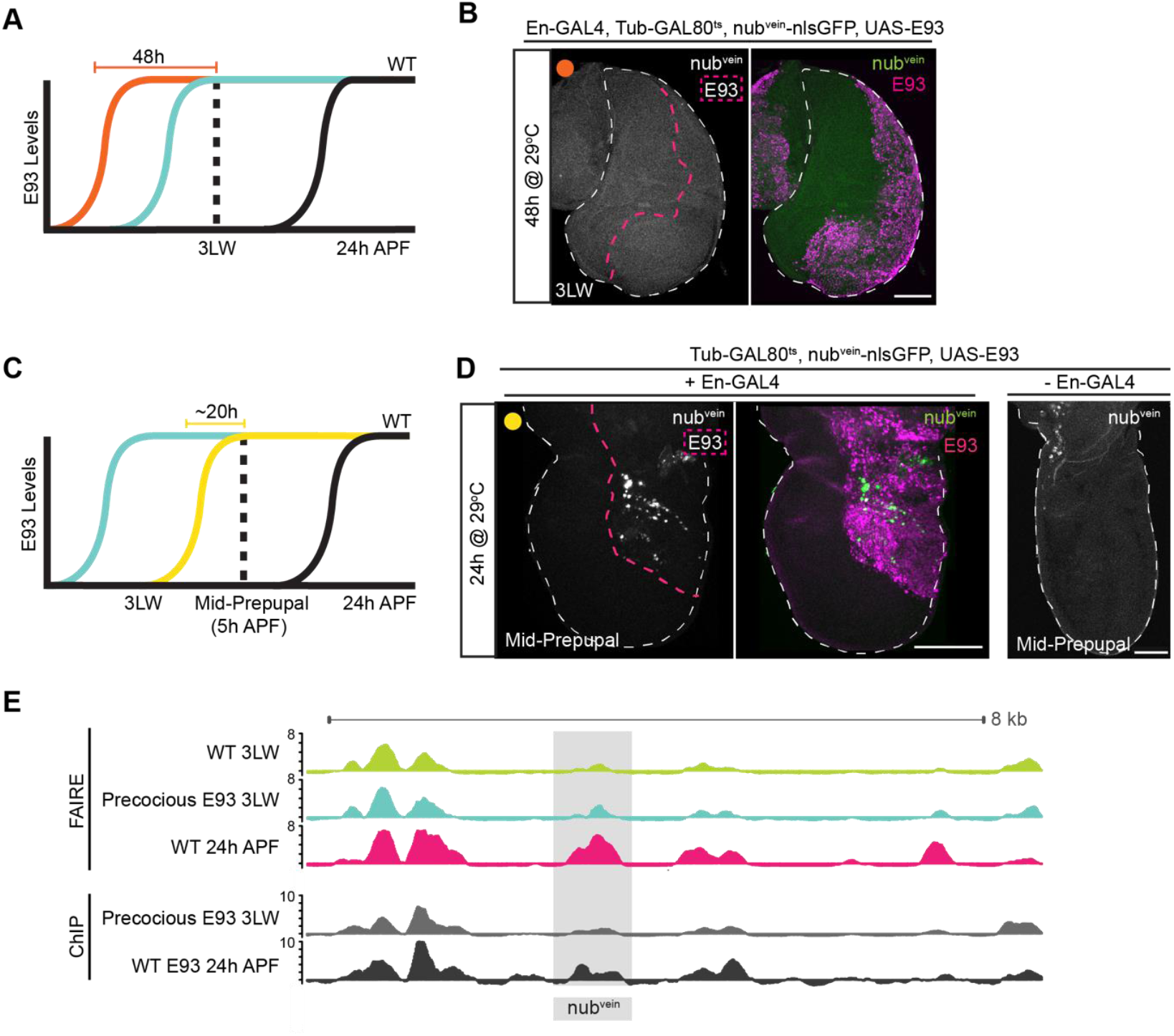
*Nub^vein^* acquires competence to respond to precocious E93 by mid-prepupal stage. (A) Schematic depicting the timing of prolonged E93 induction (orange) relative to other precocious E93-induction experiments (teal) and to wild-type E93 expression (black). (B) Confocal image of *nub*^*vein*^ activity (green) in a 3LW wing disc in which E93 (magenta) was precociously expressed for 48-hours. (C) Schematic depicting the timing of E93 induction for mid-prepupal wings (yellow) relative to other precocious E93-induction experiments (teal) and to wild-type E93 expression (black). (D) Confocal images of *nub*^*vein*^ activity (green) in mid-prepupal wings in which E93 (magenta) was precociously expressed for 20-hours. A control wing lacking *En-GAL4* is shown at the right. (E) Browser shot showing FAIRE-seq and ChIP-seq signal (z-scores) at the *nub*^*vein*^ enhancer (shaded region). Immunofluorescence scale bars = 100µm. Numbers on browser scale bars denote scale maximum.

The presence of ectopic and orphan binding sites suggests E93 binds to these sites in a context-specific manner. We therefore sought to define features of these sites that may explain the observed differences in endogenous and precocious E93 binding. We find that precocious E93 exhibits increased ChIP-seq signal at entopic sites relative to ectopic sites in 3LW wing imaginal discs (**Figure 3C**). Similarly, endogenous E93 exhibits decreased signal at orphan sites relative to entopic sites. These findings suggest that E93 has reduced ability to bind both ectopic and orphan sites relative to entopic sites (**Figure 3C**). However, examination of the DNA sequence at E93 binding sites revealed no differences in E93 motif enrichment or motif quality relative to entopic sites (**Figure 3D,E**). Therefore, the determinants that confer temporal-specific binding to orphan sites or that permit precocious E93 binding to ectopic sites remain unclear.

### Precocious E93 expression is sufficient to regulate chromatin accessibility

The ChIP-seq data described above demonstrate that the majority of targets bound by endogenous E93 in late wings are also bound by precocious E93 in early wings. We next sought to determine the impact of precocious E93 binding on chromatin accessibility. Our prior findings from E93 mutant wings suggested that E93 may function as a competence factor by controlling chromatin accessibility at target enhancers. To further test this model, we generated genome-wide open chromatin profiles by performing FAIRE-seq in wing imaginal discs precociously expressing E93. Comparison of these profiles with wild-type wing imaginal disc FAIRE-seq profiles revealed extensive changes in open chromatin. Using conservative thresholds to define differentially-accessible sites bound by E93 (DESeq2 adjusted p value < 0.05 and log_2_ fold change ≥ 1), we identified 282 sites that decrease in accessibility, 846 sites that increase in accessibility, and 7,376 sites which remain static in response to precocious E93 expression (**Figure 4 A-C**). Notably, the ratio of sites that open relative to those that close in precocious E93 early wings is similar to the ratio of sites that depend on E93 for opening and closing in wild-type late wings we previously identified in E93 loss-of-function FAIRE-seq experiments (Uyehara et al., 2017). This indicates that the ability of precociously-expressed E93 to open chromatin relative to its ability to close chromatin is similar to the abilities of endogenous E93 to regulate chromatin accessibility, despite being expressed outside of its normal developmental context.

To determine the impact of the observed changes in chromatin accessibility induced by precocious E93 expression on transcriptional regulation, we examined FAIRE-seq profiles at the E93 target enhancers described above. Accessibility of the *br*^*disc*^ enhancer strongly decreases in precocious E93 wing discs (**Figure 4D**), consistent with its deactivation by precocious E93 in transgenic reporter assays. The *br*^*disc*^ enhancer normally closes between L3 and 24-hour pupae, raising the question as to whether any of the other 281 sites that decrease in accessibility in response to precocious E93 expression also close over time in wild-type wings. Remarkably, 95% of sites that decrease in accessibility in precocious E93 wing discs also decrease in accessibility during wild-type development (**Figure 4G**), suggesting that precocious E93 expression recapitulates the normal sequence of enhancer closing. Examination of FAIRE-seq profiles at the *tnc* enhancers revealed changes in chromatin accessibility that were also consistent with the effects of precocious E93 expression on transgenic reporter activity. *Tnc*^*wv*^ and to a lesser extent *tnc*^*blade*^ increase in accessibility in response to precocious E93 (**Figure 4E**), consistent with the activation of both enhancers in transgenic reporter assays. At the genome-wide level, 73% of the sites that increase in response to precocious E93 expression also increase in accessibility during wild-type development (**Figure 4H**). Thus, the directionality of chromatin accessibility changes is preserved in wings precociously expressing E93 relative to the normal sequential changes in accessibility that occur in wild-type wings. Thus, for a subset of its target sites, E93 expression functions as an instructive cue that triggers a response in enhancer accessibility. However, E93 expression does not determine the directionality of this response. Instead, the concordance in directionality of accessibility changes between precocious E93 and wild-type development suggests that the main determinant controlling the opposing responses of different target enhancers to E93 expression is the DNA sequence of the enhancers.

### Developmental context informs the response of *nub^vein^* to precocious E93

While approximately 1,100 sites change in accessibility in response to precocious E93 expression, the majority of sites bound by precocious E93 do not change in accessibility in early wings although many are dynamic during wild-type development (**Figure 4C,F,I**). Similarly, the *nub*^*vein*^ enhancer depends on E93 for opening in pupal wings, but it fails to activate or open in response to precocious E93 expression in early wings (**Figure 2G**, **5E**, Uyehara et al., 2017). We considered the possibility that *nub*^*vein*^ requires prolonged E93 exposure, relative to E93-sensitive enhancers such as *tnc*^*blade*^, in order to become responsive, reasoning that prolonged exposure might allow time for E93-initiated indirect effects, such as induction of a co-regulator. To test this hypothesis, we doubled the duration of *nub*^*vein*^ exposure to E93 (from 24-hours to 48-hours) by inducing E93 expression earlier in development before dissecting at the same developmental stage as the experiments described above (3LW) (**Figure 5A**). Despite the prolonged exposure to E93, we still observed no change in the activity of the *nub*^*vein*^ reporter (**Figure 5B**). We next examined the possibility that E93 may require additional developmental inputs in order to activate the *nub*^*vein*^ enhancer. To test this hypothesis, we precociously expressed E93 for the same duration as in our initial experiments (20-hours), but instead of dissecting at 3LW, we dissected at a later developmental stage (mid-prepupal wings at 5hAPF, approximately 12-hours later than our initial experiments). Using this experimental design, we detected clear activation of the *nub*^*vein*^ reporter in a subset of E93-expressing cells (**Figure 5D**). Thus, the ability of the *nub*^*vein*^ enhancer to respond to precocious E93 is dependent on developmental context. It does not respond to E93 in third instar larvae regardless of the duration of E93 expression. However, it does respond to E93 in prepupal wings, suggesting a change in the regulatory environment during the larval-to-prepupal transition that makes *nub*^*vein*^ competent to respond to E93.

### Temporal dynamics of chromatin accessibility predict a context-dependent role for E93

The findings described above indicate that precocious E93 expression is sufficient to control accessibility and activity of some target sites, but other targets require additional developmentally-regulated inputs in order to respond to E93. To gain insight into the extent to which developmental context influences E93 target site responsivity, we examined the timing of chromatin accessibility changes in wild-type wings. K-means clustering of FAIRE-seq data for E93-bound sites across six time points in wild-type wing development revealed eight distinct temporal chromatin accessibility profiles (**Figure 6A**). Notably, the *br*^*disc*^, *nub*^*vein*^, and *tnc* enhancers fall into different clusters. *br*^*disc*^ falls into cluster 2 with other E93 targets that close between 6 and 18-hours hAPF (**Figure 6B**). The *tnc*^*blade*^ and *tnc*^*wv*^ enhancers fall into cluster 3 with other E93 targets that open between 6 and 18hAPF (**Figure 6B**). Finally, the *nub*^*vein*^ enhancer falls into cluster 5 with other E93 targets that open even later in pupal wing development (**Figure 6B,** Uyehara et al., 2017). Since each of these enhancers are *bona fide* E93 targets, their separation into different clusters suggests that E93 regulates target enhancers over a relatively wide range of prepupal and pupal wing development. Consistent with this interpretation, western blotting of wild-type wings at six-hour intervals surrounding the larval-to-pupal transition demonstrate that E93 expression overlaps the time points that exhibit changes in chromatin accessibility (**Figure 6C**). These findings indicate that E93 functions over a broad window of development to control enhancer activity and accessibility, and that this broad window is subdivided into narrower windows through interactions with other developmentally-regulated factors.

**Fig. 6.**
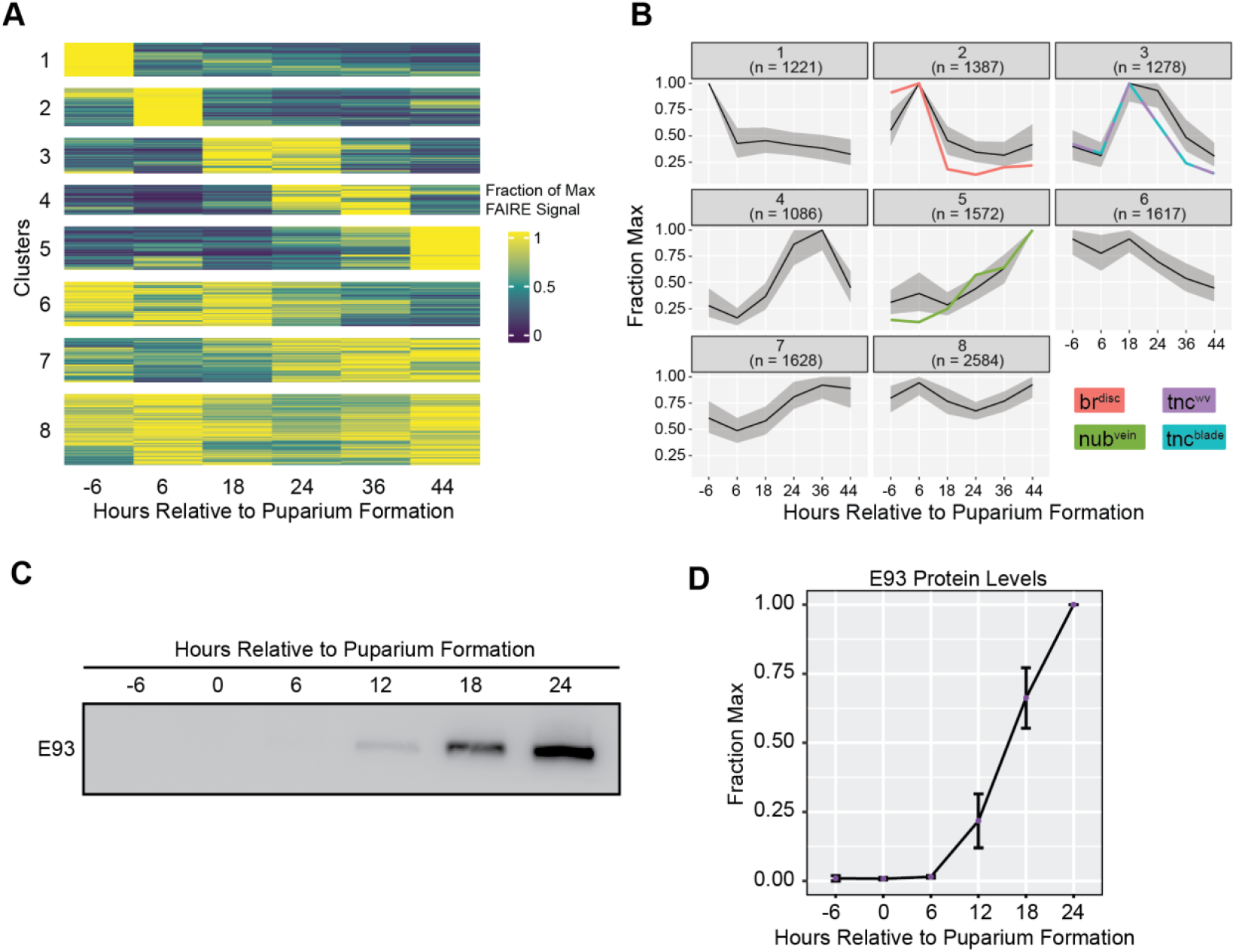
Chromatin accessibility at E93 targets changes over a broad developmental window. (A) Heatmap of chromatin accessibility over time within E93 ChIP peaks represented as fraction of maximum accessibility clustered using k-means (k=8). (B) Line plots depicting FAIRE signal for each cluster in A. Black lines represent the median fraction of max FAIRE signal and grey area represents the interquartile range. Where indicated, accessibility within ChIP-peaks that overlap enhancers are plotted in color indicated by inset labels. (C) Western blot of E93 levels in WT wings over time. (D) Quantification of E93 protein levels as measured by n = 3 western blots, normalized to total protein. Error bars = SE of Mean.

## DISCUSSION

### E93 defines a broad temporal window that is further subdivided by additional temporal cues

Despite its ability to regulate chromatin accessibility at a subset of target enhancers in wing imaginal discs, precocious E93 expression is insufficient to control accessibility at many other sites to which it binds. This suggests that, like many transcription factors, E93 function can also depend on context. Our examination of the *nub*^*vein*^ enhancer provides potential mechanistic insight to this context-specific activity. Although *nub*^*vein*^ depends on E93 for opening and activation in wild-type pupal wings, precocious E93 expression in larval wing discs does not result in *nub*^*vein*^ activation or in increased chromatin accessibility. *Nub*^*vein*^ remains inactive even with prolonged exposure to E93 at larval stages, suggesting that its activation is not dependent on a downstream effector of E93 activity. Instead, *nub*^*vein*^ exhibits precocious activity in response to E93 expression only after progression through the larval-to-prepupal transition. This switch in responsivity of *nub*^*vein*^ as a function of developmental stage rather than duration of E93 exposure indicates that there is a change in the *trans*-regulatory environment that is independent of E93 activity. This change could be either a developmentally-programmed gain of an activator that works in conjunction with E93 or the loss of a repressor that blocks E93.

Considering this switch in *nub*^*vein*^ responsivity in combination with the observation that E93 target sites exhibit changes in chromatin accessibility over two days during metamorphosis helps to inform a model for temporal gene regulation genome-wide (**Figure 7**). We propose that expression of E93 defines a broad temporal window of pupal identity that is further subdivided by the activity of parallel developmentally-regulated pathways. One potential pathway that could provide an additional source of temporal information is the ecdysone pathway. In addition to E93 induction, ecdysone signaling initiates expression of multiple other transcription factors that could work in conjunction with E93 or antagonize its activity. For example, the ecdysone-induced transcription factors *broad* and *ftz-f1* are both dynamically expressed surrounding the larval-to-prepupal transition (Broadus et al., 1999; Crossgrove et al., 1996; Karim et al., 1993). In addition, the levels of circulating ecdysone could impact E93 function. E93 binds to the ecdysone hormone receptor, EcR/Usp, through its LXXLL nuclear receptor interaction motif (Liu et al., 2015). As hormone levels rise and fall during the ecdysone pulses that trigger the larval-to-prepupal and prepupal-to-pupal transitions, E93/EcR/Usp complexes may assemble on target enhancers in a hormone-dependent manner (Uyehara and McKay, 2019; Yao et al., 1993). For example, it is possible that high ecdysone titers prevent E93 binding to the *nub*^*vein*^ enhancer in larval wing discs, and the decrease in these titers in prepupal wings enables E93 binding. Finally, developmentally-programmed changes in the expression patterns of spatial transcription factors that work with E93 could also further subdivide temporal windows of E93 action on target genes.

**Fig. 7.**
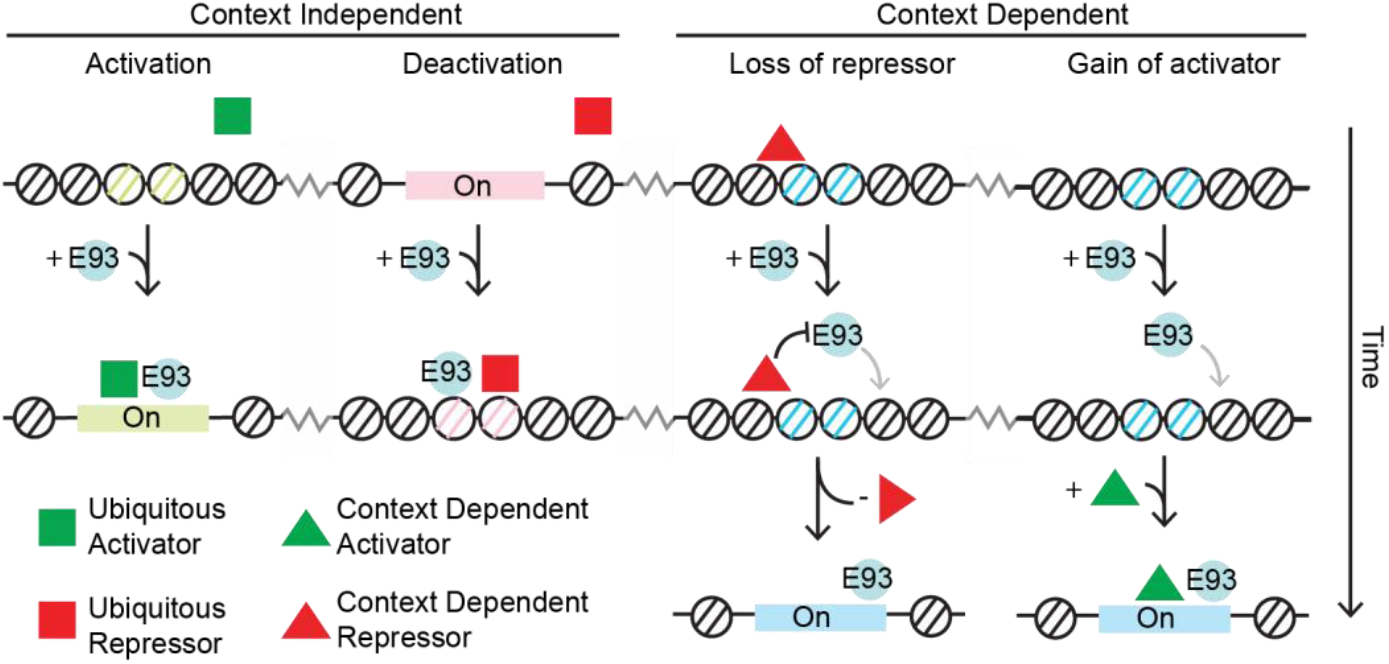
Models of E93 function at target enhancers. E93 expression is sufficient to bind and regulate chromatin accessibility at a subset of its target enhancers. However, other target enhancers respond to E93 only in certain developmental contexts. This leads to a model wherein E93 works with co-regulatory proteins that are either ubiquitously-expressed (squares) or developmentally-regulated (triangles). These factors work with E93 as activators (green) or repressors (red) at target enhancers. Note that context-dependent deactivation is a possible outcome that is not shown.

### Temporal transcription factors as determinants of developmental competence

Spatial cues are iteratively used during development to produce distinct transcriptional outcomes. Many of these spatial inputs come in the form of transcription factors that are expressed at multiple stages of development. However, it is unclear why these factors regulate their given targets only at select times. The work presented in this study indicate that E93 expression provides competence for target enhancers to respond to spatially-restricted inputs. Precocious expression of E93 activates the *tnc*^*blade*^ enhancer in the proximal hinge of wing imaginal discs, similar to its wild-type pattern of activity in pupal wings. Likewise, the *tnc*^*wv*^ enhancer is precociously activated in cells of the pouch near to high levels of Dpp signaling, similar to the cells that activate *tnc*^*wv*^ later in pupal wings. Notably, neither of these enhancers become active in all cells that precociously express E93. Instead, precocious E93 expression activates these enhancers only in populations of cells similar to those in which they become active later in development. This suggests that provision of E93 enables these enhancers to respond to the spatial cues that regulate their activity later. Consistent with this interpretation, precocious E93 expression in developing legs provides competence to the *Distal-less* gene to respond to EGFR signaling specifically in bract cells (Mou et al., 2012). Importantly, like the spatial cues that regulate the *tnc* enhancers, the EGFR pathway is active in both early and late legs, and yet EGFR is only capable of activating *Distal-less* in the presence of E93. Thus, the spatial cues present prior to E93 expression are insufficient to activate the *Distal-less* gene or the E93-target enhancers described here, indicating that E93 is the key determinant for unlocking their activities. Our ChIP-seq data demonstrate that the effects of E93 on target enhancers are direct. More importantly, FAIRE-seq reveals that the mechanism by which E93 provides competence to target enhancers lies at least in part by controlling chromatin accessibility. Together, these findings are consistent with a model in which E93 functions as a temporal cue by controlling the chromatin accessibility landscape. At a subset of enhancers, E93 binding results in chromatin opening, which enables binding of other transcription factors that control the spatial pattern of enhancer activity. At a different subset of enhancers that are already accessible, E93 binding results in chromatin closing, making these enhancers refractory to transcription factor binding and enabling redeployment of these spatial inputs to other targets. Together, these activities help to control the sequence of gene expression changes that drive development forward in time.

### The importance of target DNA sequence in determining context-specific transcription factor function

While it is clear that transcription factors often contain both activating and repressing roles during development, the determinants of this context-specific function are poorly understood. In this study, we find that E93 expression both activates and represses different target enhancers in the same cells at the same time. The *br*^*disc*^ enhancer is expressed across the entire larval wing disc and is both closed and deactivated in response to precocious E93 expression. Conversely, the *tnc*^*blade*^ and *tnc*^*wv*^ enhancers both opened and activated in response to precocious E93 expression. The pattern of *br*^*disc*^ overlaps the precocious expression pattern of both *tnc* enhancers, thus strongly indicating that E93 expression is sufficient to enact two opposing functions (activation and deactivation) simultaneously during development. Genomic profiling of the chromatin accessibility landscape suggests that E93 acts at hundreds of sites across the genome to both open and close chromatin when precociously expressed. This indicates that the ability of E93 to increase and decrease accessibility of target enhancers is not exclusively due to stage-specific expression of co-regulators or a temporally-regulated modification of E93 to make it a dedicated activator or repressor. Instead, the observation that sites that open or close in response to precocious E93 mirror changes in the wild-type accessibility program indicates that the DNA sequence of target enhancers encodes the response to E93. Therefore, the key determinants of E93 function are likely the transcription factors that co-occupy target enhancers with E93.

### E93 expression is sufficient for changes in chromatin accessibility

Our prior work demonstrated that E93 is required for proper chromatin accessibility at temporally-dynamic enhancers during pupal wing development. However, it remained unclear whether E93 initiates these changes. An alternative possibility was that E93 maintains a chromatin landscape established by other factors. The findings presented here reveal that precocious E93 expression is sufficient to induce changes in chromatin accessibility, indicating that E93 initiates opening and closing of at least a subset of its target enhancers during wild-type development. The sufficiency of E93 to regulate chromatin accessibility might mean that E93 works on its own. However, our findings that E93 responsiveness also depends on developmental context indicates that E93 likely works with other co-regulators to control nucleosome occupancy at target sites (**Figure 7**). Identifying these co-regulatory factors will be important for understanding the mechanisms by which temporal transcription factors drive the sequence of gene expression changes that underlie progressive specification of cell fates in development.

## METHODS

### *Drosophila* culture and genetics

Either *Ci-GAL4* or *En-GAL4* lines were used for enhancer reporter experiments. Crosses were raised at room temperature (22°C) and vials were shifted to 29°C for 22–33-hours to induce E93 expression, unless otherwise indicated. For the experiments presented in Figure 5B, vials were shifted to 29°C for 48-hours. Wandering third instar larvae were collected for all experiments, except for the experiments presented in Figure 5D, in which mid-prepupae (5hAPF) were collected after 20-hours of E93 induction. The *vg-GAL4, tub*>*CD2*>*GAL4, UAS-GFP, UAS-FLP; GAL80*^*ts*^ driver was used for genomics experiments. Embryos were collected for 6-hours on apple plates at 25°C and then maintained at 25°C for an additional 24-hours. 24–30-hours GFP-positive larvae were picked, transferred to vials, and raised for 12-hours at 29°C to induce flip-out of the CD2 cassette. Vials were transferred to 18°C for 4.5 days, then switched back to 29°C for 15-hours to induce E93 expression. Wing discs from wandering third instar larvae were dissected for FAIRE and ChIP experiments.

List of fly stocks used:

*w; Tub-GAL80*^*ts*^*; tm2/tm6b* (BDSC 7108)

*yw;; UAS-E93-3xHA* (FlyORF F000587, Bischof et al., 2013)

*yw; vg-GAL4, UAS-FLP, UAS-GFP, Tub*>>*CD2*>>*EGAL4 / CyO* (Crickmore and Mann, 2006)

*yw; En-GAL4;* (Gift of Greg Matera)

*yw; Ci-GAL4;* (Gift of Robert Duronio)

*yw;; broad*^*disc*^*-tdTomato* (Uyehara et al., 2017)

*yw;; nub*^*vein*^*-nlsGFP* (Uyehara et al., 2017)

*yw;; tnc*^*wv*^*-tdTomato* (Uyehara et al., 2017)

*yw;; tnc*^*blade*^*-tdTomato* (Uyehara et al., 2017)

Details on genotypes for particular experiments are available upon request.

### Immunofluorescence and Image Analysis

Immunofluorescence experiments and confocal imaging were performed as previously described (McKay and Lieb, 2013). The following antibodies were used: rabbit anti-E93 (1:2500, Uyehara et al., 2017), mouse anti-HA (1:1000, Sigma H3663), guinea pig anti-Teashirt (1:1000, Zirin and Mann, 2007), mouse anti-Delta (1:350, DSHB C594.9b), mouse anti-Wingless (1:25, DSHB 4D4), goat anti-rabbit Alexa 633 (1:1000, Invitrogen A21071), goat anti-rabbit Alexa 594 (1:1000, Invitrogen A111037), goat anti-mouse Alexa 633 (1:1000, Invitrogen A21052), goat anti-mouse Alexa 594 (1:1000, Invitrogen A11032), and goat anti-guinea pig Alexa 647 (1:1000, Invitrogen A21450). Precocious expression and WT levels of E93 at ~30h APF were quantified by immunofluorescence using the rabbit anti-E93 antibody and the goat anti-rabbit Alexa 633 secondary. Fluorescent intensity was measured using ImageJ (Schindelin et al., 2012). Late pupal wings with WT E93 were combined with 3LW imaginal wing discs precociously expressing E93 (using either the Ci-GAL4 or Vg-GAL4 driver) in the same tube for incubation with primary and secondary antibodies. Early and late wings were then mounted on the same slide and imaged with identical settings (Leica Confocal SP5). E93 levels were quantified by measuring mean grey value in 25×25 pixel selections (10 selections per wing and 3 wings each for early and late time points). E93 signal was normalized to background by dividing each E93 measurement by the mean background, which was calculated from 9 25×25 pixel selections in E93-negative portions of tissue in each experiment.

### High throughput sequencing & data analysis

FAIRE-seq and ChIP-seq were performed as previously described (Uyehara 2017). Briefly, ChIP experiments were performed in duplicate using a minimum of 200 wings for each replicate. Control genotypes contained the *GAL4* driver but lacked the *UAS-E93-3xHA* transgene. Immunoprecipitation was performed using 5 µg of Rabbit anti-HA (Abcam, ab9110). FAIRE-seq in precocious E93-expressing wings was performed using between 45 and 60 wings in duplicate. Sequencing reads were aligned to the dm3 reference genome with Bowtie2 (Langmead and Salzberg, 2012). ChIP peaks were called with MACS2 (Zhang et al., 2008) on each replicate using as background reads from the control genotype (precocious E93 experiments) or from a sonicated genomic DNA library (wild-type 24hAPF E93). ChIP peaks that overlapped between biological replicates were used for analysis. E93 binding categories were identified by intersecting the resulting peak lists from precocious E93 ChIP and wild-type E93 ChIP using the ChIPpeakAnno package from bioconductor (Gentleman et al., 2004; R Core Team, 2017). Summits from the resulting union ChIP peak list were recomputed from aligned reads from pooled replicates using FunChip (setting d = 125) (v1.0.0) (Parodi et al., 2017). Summits from entopic and orphan sites were computed from wild-type late E93 ChIP-seq, while summits from ectopic sites were computed from precocious E93 ChIP. Chromatin accessibility differences within precocious E93 ChIP peaks were identified by counting FAIRE-seq reads within the union set of E93 ChIP peaks using featureCounts (setting allowMultioverlap = T) from Rsubread and testing for differential accessibility with DESeq2 using an adjusted p value < 0.05 and an absolute log2FoldChange > 1 (Liao et al., 2019; Love et al., 2014). Concordance of precocious chromatin accessibility changes with wild-type chromatin accessibility changes was determined using DESeq2, using an adjusted p value < 0.05. Average signal line plots were generated using seqplots and ggplot2 from z-score normalized bigwig files at 10 base-pair resolution (Stempor and Ahringer, 2016; Wickham, 2009). Signal tracks were rendered in R with Gviz and cowplot (Hahne and Ivanek, 2016; Wilke, 2017).

### Motif Scanning

The dm3 assembly of the *Drosophila melanogaster* genome was scanned for the E93 motif from the FlyFactor Survey database using FIMO v4.12.0 (setting –thresh 0.01 –max-strand –text –skip-matchd-sequence) (Grant et al., 2011; Zhu et al., 2011). Motifs overlapping a 20-base pair window around ChIP peak summits were identified using GenomicRanges findOverlaps (Lawrence et al., 2013). Motif number per window was quantified by directly counting these overlaps. For each peak category, motif quality within these windows was compared by using the ‘oneway.test’ function in R.

### Western Blotting

Wing discs were dissected from *E93*^*GFSTF*^ animals at 6-hour intervals relative to puparium formation by staging animals as white prepupae (third-instar wandering larvae were used as the –6h timepoint). Western blots were performed as previously described, with the following changes. 20 wings were collected per timepoint in phosphate-buffered saline (PBS) in 1.5 mL microcenterfuge tubes (Leatham-Jensen et al., 2019). Samples were subsequently spun at 1000 rcf × 5 minutes to pellet wings, PBS was removed and samples were stored at −80 °C. Samples were prepared by heating at 95 °C for 10 minutes in 45 µL of hot Laemmli SDS-PAGE loading buffer. Samples were loaded onto a 7.5% Biorad stain-free TGX gel (Product #4568024) and run for 60 minutes at 100 V. Total protein stains were collected by laying the PAGE gel directly onto a UV transilluminator for 3 minutes and imaged on an Amersham Imager 600; the gel was kept hydrated with distilled water during all total protein crosslinking and imaging steps. After imaging the total protein stain, protein was transferred to a 0.2 µm nitrocellulose membrane at 100 V for 60 minutes. E93^GFSTF^ protein was detected using rabbit-anti-GFP (1:5000, abcam ab290), HRP-conjugated secondary (1:10000 donkey anti-Rabbit-HRP, GE Healthcare #NA934V) and Amersham ECL prime detection kit (GE healthcare, RPN2232). Blots were imaged on an Amersham Imager 600. Blots and total-protein signal were quantified with FIJI, where individual replicates were scaled first to total protein then relative to the timepoint with maximum E93 signal (in all cases, 24h APF). Signal quantification represents three independent western blots.

---

**Fig. S1.**
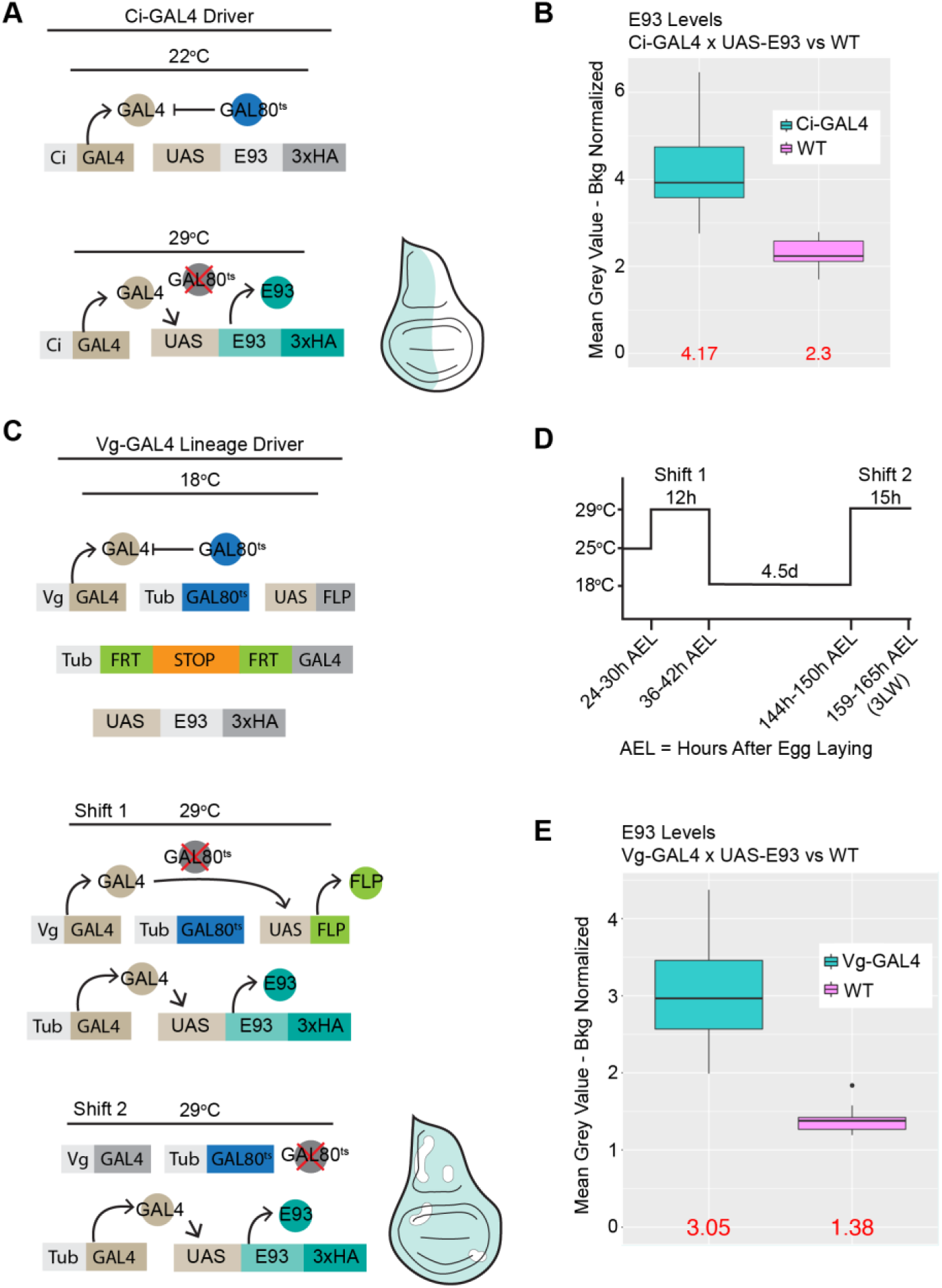
Ectopic E93 expression system leads to two-fold higher levels of E93 relative to wild type. (A) Schematic of molecular details of *Ci-GAL4/GAL80*^ts^ control of the *UAS-E93-3xHA* transgene. (B) Boxplots showing quantification of E93 levels in 3LW discs using the Ci-GAL4 driver relative to wild-type (WT) E93 levels at 30h APF. (D) Schematic of molecular details of the *Vg-GAL4* lineage tracing diver. Shift 1 removes the STOP cassette in first instar wings. Shift 2 induces E93 expression in third instar wings. Due to the efficiency of flip-out some portion of each disc remains WT (white regions). (D) Temperature shifting scheme for *Vg-GAL4* driver with lineage tracking. (E) Boxplots showing quantification of E93 levels in 3LW discs using the *Vg-GAL4* lineage tracing driver relative to WT. Averages indicated in red. n = 30 (10 measurements across 3 wings) per condition.

**Fig. S2.**
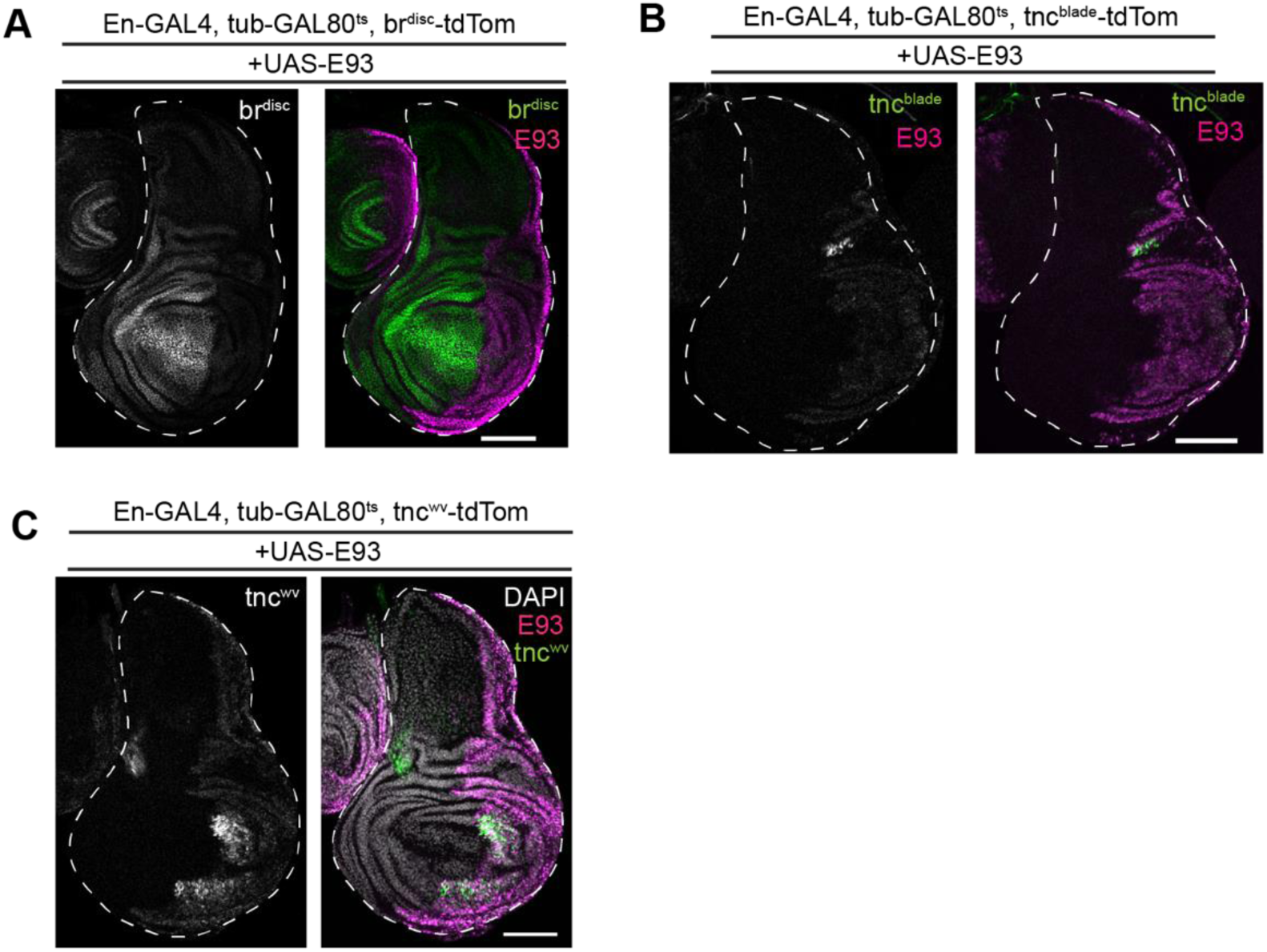
Patterns of enhancer activity with precocious E93 in posterior compartment of wing imaginal disc. (A) Confocal image of *br*^disc^ activity (green) in a wing disc precociously-expressing E93 (magenta) using the *En-GAL4* driver. (B) Confocal image of *tnc*^blade^ activity (green) in a wing disc precociously-expressing E93 (magenta) using the *En-GAL4* driver. (C) Confocal image of *tnc*^wv^ activity (green) in a wing disc precociously-expressing E93 (magenta) using the *En-GAL4* driver. Scale bars = 100μm.

## REFERENCES

Baehrecke, E. H. and Thummel, C. S. (1995). The Drosophila E93 Gene from the 93F Early Puff Displays Stage- and Tissue-Specific Regulation by 20-Hydroxyecdysone. Dev. Biol. 171, 85–97.

Bischof, J., Bjorklund, M., Furger, E., Schertel, C., Taipale, J. and Basler, K. (2013). A versatile platform for creating a comprehensive UAS-ORFeome library in Drosophila. Development 140, 2434–2442.

Broadus, J., McCabe, J. R., Endrizzi, B., Thummel, C. S. and Woodard, C. T. (1999). The Drosophila beta FTZ-F1 orphan nuclear receptor provides competence for stage-specific responses to the steroid hormone ecdysone. Mol. Cell 3, 143–9.

Crickmore, M. A. and Mann, R. S. (2006). Hox Control of Organ Size by Regulation of Morphogen Production and Mobility. Science (80-.). 313, 63–68.

Crossgrove, K., Bayer, C. A., Fristrom, J. W. and Guild, G. M. (1996). The Drosophila Broad-Complex early gene directly regulates late gene transcription during the ecdysone-induced puffing cascade. Dev. Biol. 180, 745–58.

De Celis, J. F. (1997). Expression and function of decapentaplegic and thick veins during the differentiation of the veins in the Drosophila wing. Development 124, 1007–18.

Doe, C. Q. (2017). Temporal Patterning in the *Drosophila* CNS. Annu. Rev. Cell Dev. Biol. 33, 219–240.

Fraichard, S., Bougé, A.-L., Kendall, T., Chauvel, I., Bouhin, H. and Bunch, T. A. (2010). Tenectin is a novel αPS2βPS integrin ligand required for wing morphogenesis and male genital looping in Drosophila. Dev. Biol. 340, 504–517.

Gentleman, R. C., Carey, V. J., Bates, D. M., Bolstad, B., Dettling, M., Dudoit, S., Ellis, B., Gautier, L., Ge, Y., Gentry, J., et al. (2004). Bioconductor: open software development for computational biology and bioinformatics. Genome Biol. 5, R80.

Grant, C. E., Bailey, T. L. and Noble, W. S. (2011). FIMO: scanning for occurrences of a given motif. Bioinformatics 27, 1017–1018.

Guo, Y., Flegel, K., Kumar, J., McKay, D. J. and Buttitta, L. A. (2016). Ecdysone signaling induces two phases of cell cycle exit in Drosophila cells. Biol. Open 5, 1648–1661.

Hahne, F. and Ivanek, R. (2016). Visualizing Genomic Data Using Gviz and Bioconductor. In Methods in molecular biology (Clifton, N.J.), pp. 335–351.

Holguera, I. and Desplan, C. (2018). Neuronal specification in space and time. Science (80-.). 362, 176–180.

Karim, F. D., Guild, G. M. and Thummel, C. S. (1993). The Drosophila Broad-Complex plays a key role in controlling ecdysone-regulated gene expression at the onset of metamorphosis. Development 118, 977–88.

Kumar, N., Tsai, Y.-H., Chen, L., Zhou, A., Banerjee, K. K., Saxena, M., Huang, S., Toke, N. H., Xing, J., Shivdasani, R. A., et al. (2019). The lineage-specific transcription factor CDX2 navigates dynamic chromatin to control distinct stages of intestine development. Development 146, dev172189.

Langmead, B. and Salzberg, S. L. (2012). Fast gapped-read alignment with Bowtie 2. Nat. Methods 9, 357–359.

Lawrence, M., Huber, W., Pagès, H., Aboyoun, P., Carlson, M., Gentleman, R., Morgan, M. T. and Carey, V. J. (2013). Software for Computing and Annotating Genomic Ranges. PLoS Comput. Biol. 9, e1003118.

Leatham-Jensen, M., Uyehara, C. M., Strahl, B. D., Matera, A. G., Duronio, R. J. and McKay, D. J. (2019). Lysine 27 of replication-independent histone H3.3 is required for Polycomb target gene silencing but not for gene activation. PLOS Genet. 15, e1007932.

Liu, X., Dai, F., Guo, E., Li, K., Ma, L., Tian, L., Cao, Y., Zhang, G., Palli, S. R. and Li, S. (2015). 20-Hydroxyecdysone (20E) Primary Response Gene E93 Modulates 20E Signaling to Promote Bombyx Larval-Pupal Metamorphosis. J. Biol. Chem. 290, 27370–83.

Martín-Blanco, E., Roch, F., Noll, E., Baonza, A., Duffy, J. B. and Perrimon, N. (1999). A temporal switch in DER signaling controls the specification and differentiation of veins and interveins in the Drosophila wing. Development 126, 5739–47.

McKay, D. J. and Lieb, J. D. (2013). A common set of DNA regulatory elements shapes Drosophila appendages. Dev. Cell 27, 306–18.

Mou, X., Duncan, D. M., Baehrecke, E. H. and Duncan, I. (2012). Control of target gene specificity during metamorphosis by the steroid response gene E93. Proc. Natl. Acad. Sci. U. S. A. 109, 2949–54.

O’Keefe, D. D., Thomas, S., Edgar, B. A. and Buttitta, L. (2014). Temporal regulation of Dpp signaling output in the *Drosophila* wing. Dev. Dyn. 243, 818–832.

Parodi, A. C. L., Sangalli, L. M., Vantini, S., Amati, B., Secchi, P. and Morelli, M. J. (2017). FunChIP: an R/Bioconductor package for functional classification of ChIP-seq shapes. Bioinformatics 33, 2570–2572.

Pasquinelli, A. E. and Ruvkun, G. (2002). Control of Developmental Timing by MicroRNAs and Their Targets. Annu. Rev. Cell Dev. Biol. 18, 495–513.

Pavlopoulos, A. and Akam, M. (2011). Hox gene Ultrabithorax regulates distinct sets of target genes at successive stages of Drosophila haltere morphogenesis. Proc. Natl. Acad. Sci. 108, 2855–2860.

R Core Team (2017). R: A Language and Environment for Statistical Computing.

Schindelin, J., Arganda-Carreras, I., Frise, E., Kaynig, V., Longair, M., Pietzsch, T., Preibisch, S., Rueden, C., Saalfeld, S., Schmid, B., et al. (2012). Fiji: an open-source platform for biological-image analysis. Nat. Methods 9, 676–682.

Sen, S. Q., Chanchani, S., Southall, T. D. and Doe, C. Q. (2019). Neuroblast-specific open chromatin allows the temporal transcription factor, Hunchback, to bind neuroblast-specific loci. Elife 8,.

Stempor, P. and Ahringer, J. (2016). SeqPlots - Interactive software for exploratory data analyses, pattern discovery and visualization in genomics. Wellcome Open Res. 1, 14.

Uyehara, C. M. and McKay, D. J. (2019). Direct and widespread role for the nuclear receptor EcR in mediating the response to ecdysone in Drosophila. Proc. Natl. Acad. Sci. U. S. A. 116, 9893–9902.

Uyehara, C. M., Nystrom, S. L., Niederhuber, M. J., Leatham-Jensen, M., Ma, Y., Buttitta, L. A. and McKay, D. J. (2017). Hormone-dependent control of developmental timing through regulation of chromatin accessibility. Genes Dev. 31, 862–875.

Wickham, H. (2009). ggplot2?: elegant graphics for data analysis. Springer.

Wilke, C. O. (2017). cowplot: Streamlined Plot Theme and Plot Annotations for “ggplot2”. R package version 0.9.4. https://CRAN.R-project.org/package=cowplot. R Packag. version0.9.2 https://cran.r-project.org/package=cowplot.

Yao, T.-P., Forman, B. M., Jiang, Z., Cherbas, L., Chen, J.-D., McKeown, M., Cherbas, P. and Evans, R. M. (1993). Functional ecdysone receptor is the product of EcR and Ultraspiracle genes. Nature 366, 476–479.

Zhang, Y., Liu, T., Meyer, C. A., Eeckhoute, J., Johnson, D. S., Bernstein, B. E., Nusbaum, C., Myers, R. M., Brown, M., Li, W., et al. (2008). Model-based analysis of ChIP-Seq (MACS). Genome Biol. 9, R137.

Zhu, L. J., Christensen, R. G., Kazemian, M., Hull, C. J., Enuameh, M. S., Basciotta, M. D., Brasefield, J. A., Zhu, C., Asriyan, Y., Lapointe, D. S., et al. (2011). FlyFactorSurvey: a database of Drosophila transcription factor binding specificities determined using the bacterial one-hybrid system. Nucleic Acids Res. 39, D111–7.

Zirin, J. D. and Mann, R. S. (2007). Nubbin and Teashirt mark barriers to clonal growth along the proximal-distal axis of the Drosophila wing. Dev. Biol. 304, 745–58.

